# Development and validation of a clinical risk score to predict the risk of SARS-CoV-2 infection from administrative data: a population-based cohort study from Italy

**DOI:** 10.1101/2020.07.23.217331

**Authors:** Valentina Orlando, Federico Rea, Laura Savaré, Ilaria Guarino, Sara Mucherino, Alessandro Perrella, Ugo Trama, Enrico Coscioni, Enrica Menditto, Giovanni Corrao

## Abstract

**Background:** The novel coronavirus (SARS-CoV-2) pandemic spread rapidly worldwide increasing exponentially in Italy. To date, there is lack of studies describing clinical characteristics of the population most at risk of infection. Hence, we aimed to identify clinical predictors of SARS-CoV-2 infection risk and to develop and validate a score predicting SARS-CoV-2 infection risk comparing it with unspecific surrogates.

**Methods:** Retrospective case/control study using administrative health-related database was carried out in Southern Italy (Campania region) among beneficiaries of Regional Health Service aged over than 30 years. For each subject with Covid-19 confirmed diagnosis (case), up to five controls were randomly matched for gender, age and municipality of residence. Odds ratios and 90% confidence intervals for associations between candidate predictors and risk of infection were estimated by means of conditional logistic regression. SARS-CoV-2 Infection Score (SIS), was developed by generating a total aggregate score obtained from assignment of a weight at each selected covariate using coefficients estimated from the model. Finally, the score was categorized by assigning increasing values from 1 to 4. SIS was validated by comparison with specific and unspecific predictors of SARS-CoV-2 infection.

**Results:** Subjects suffering from diabetes, anaemias, Parkinson’s disease, mental disorders, cardiovascular and inflammatory bowel and kidney diseases showed increased risk of SARS-CoV-2 infection. Similar estimates were recorded for men and women and younger and older than 65 years. Fifteen conditions significantly contributed to the SIS. As SIS value increases, risk progressively increases, being odds of SARS-CoV-2 infection among people with the highest SIS value (SIS=4), 1.74 times higher than those unaffected by any SIS contributing conditions (SIS=1).

**Conclusion:** This study identified conditions and diseases making individuals more vulnerable to SARS-CoV-2 infection. Our results are a decision-maker support tool for identifying population most at risk allowing adoption of preventive measures to minimize a potential new relapse damage.

## Introduction

Since December 2019, the novel coronavirus (SARS-CoV-2) pandemic spread rapidly from the Hubei province in China to 185 countries causing over 3,000,000 cases [1]. The epidemic spread to and increased exponentially in Italy, earlier than in any other western Country, having generated at the current time (June 15) over 236,000 confirmed SARS-CoV-2 infections [2].

Several hospital-based studies [3–7], including a systematic review of literature and meta-analysis [8], focused around the attempt for predicting the progression of the disease towards developing critical manifestations or death. These studies are important for the clinical practice point of view for identifying patients at whom early treatment must be guaranteed. However, as the vast majority of infections are not life-threatening [4], for the public health point of view it becomes increasingly important stratifying population for identifying people at higher risk of infection. In spite of this, at our best knowledge, no studies on this topic have been still published.

We therefore performed a large investigation based upon healthcare utilization database from the Italian Region of Campania aimed (1) to identify clinical predictors of the risk of SARS-CoV-2 infection, (2) to develop and validate a score overall predicting the risk of SARS-CoV-2 infection, and (3) to compare discriminant power of such a score with that from unspecific surrogates of clinical profile.

## Methods

### Target population and data source

Resident in Campania who were beneficiaries of the Regional Health Service, and were aged 30 years or older, formed the target population (just almost 3.9 million people, around 9% of the Italian population in that age group). Italian citizens have equal access to essential healthcare services provided by the National Health Service. The present study was carried out using information collected routinely in healthcare databases in Campania. The Campania Region Database (CaReDB) includes information on patient demographics, the electronic records of outpatient pharmacy dispensing and hospital discharges for ~6 million residents of a well-defined population in Italy (~10% of the population of Italy). CaReDB is complete and includes validated data in previous drug utilization studies [9–17]. The characteristics of CaReDB are described in S1 Table.

From the beginning of the Covid-19 epidemic, a surveillance system was implemented to collect all cases identified by reverse transcription-polymerase chain reaction (RT-PCR) testing for SARS-CoV-2. Diagnostic algorithm was based on the protocol released by the World Health Organization (WHO) [18], i.e., on nasopharyngeal swab specimens tested with at least two real-time RT PCT assays targeting different genes (E, RdRp and M) of SARS-CoV-2.

These archives abovementioned can be linked together by a unique anonymous identifier that is encrypted to protect patient privacy. Permission use anonymized data to this study was granted to the researchers of the Centro di Ricerca in Farmacoeconomia e Farmacoutilizzazione (CIRFF) by the governance board of Unità del Farmaco della Regione Campania. The research does not contain clinical studies, and all patients’ data were fully anonymized and were analysed retrospectively. For this type of study, formal consent is not required according to current national law from Italian Medicines Agency and according to the Italian Data Protection Authority, neither Ethical Committee approval nor informed consent were required for our study [19]. Our research protocol adhered to the tenets of the Declaration of Helsinki 1975 and its later amendments.

### Cases and controls

The date of Covid-19 infection diagnosis was considered as the index date and patients were extracted from the registry until June 10, 2020. A total of 4,629 subjects positive to Covid-19 were identified. Among these, we excluded i) patients with missing demographic information (N=469) and ii) patients younger than 30 years at the index date (N=663). Finally, 3,497 patients were included into the study as cases. Among them, 453 patients died during the observational period.

For each case, up to five controls were randomly selected from the target population to be matched for gender, age at index date and municipality of residence. The density incidence approach was used for selecting controls since patients who had a confirmed diagnosis of Covid-19 infection were eligible as potential controls until they became cases, and all matches had to be at risk of Covid-19 infection.

### Identifying clinical predictors of SARS-CoV-2 infection

A list of 47 diseases and conditions predictors of the risk of SARS-CoV-2 infection was carefully chosen and assessed using a modified version of the RxRiskV Index. The latter is a validated pharmaceutical-based comorbidity index derived from dispensation data using Anatomical Therapeutic Chemical (ATC) classification codes [20,21]. The RxRiskV Index was improved for our study to include updated ATC codes for medications licensed in Italy currently and adding the pertaining ICD-9 CM code for each condition. These amendments were made according to previously published works [22–29]. Individuals were classified as having one of the conditions listed if they received at least ≥2 consecutive dispensations of a drug for treatment of a specific class of disease and/or one hospital discharge with the diagnoses coded with the specific ICD-9-CM (S2 Table).

Conditional logistic regression was used to estimate odds ratios (ORs), with 90% confidence intervals (CIs), for the association between candidate predictors and the odds of SARS-CoV-2 infection. Predictors entered as dichotomous covariates into the model, i.e., with value 0 or 1 according to whether the specific condition was not or was recorded at least once within two-years prior baseline (2018-2019). Unadjusted and mutually adjusted models were fitted by including one by one covariate, and all covariates together, respectively. Power considerations suggested excluding covariates with prevalence ≤ 0.12% among controls, i.e., predictors for which our sample size was not sufficient for detecting OR of at least 3, with a 0.80 power, and by accepting a 0.10 two-sided first type error. In addition, some conditions were grouped together when strong uncertainty of algorithm did not allow for distinguishing them.

With the aim of testing the hypothesis that predictors may affect severity of clinical manifestations of SARS-CoV-2 infection, rather than infection *per se*, analyses were restricted to strata having fatal infection. Stratifications for sex and age categories (<65 years, ≥65 years) were performed as secondary analyses.

### Developing and validating a score to predict SARS-CoV-2 infection

Seven out of ten of the 3,497 1:5 case-control sets were randomly selected to form the so-called training (derivation) set. The conditional logistic regression model was fitted to compute the odds ratios as above described. The least absolute shrinkage and selection operator (LASSO) method was applied for selecting the diseases / conditions able to independently predict the SARS-CoV-2 infection [30]. The coefficients estimated from the model were used for assigning a weight at each selected covariate. In particular, a weight was assigned to each coefficient by multiplying it by 10 and rounding it to the nearest whole number [31]. The weights thus obtained were then summed to generate a total aggregate score. To simplify the system, i.e., with the aim of accounting for excessive heterogeneity of the total aggregate score, the latter was categorized by assigning increasing values of 1, 2, 3 and 4 to the categories of the aggregate score of 0, 1-2, 3-4, ≥ 5, respectively. The so obtained index was denoted SARS-CoV-2 Infection Score (SIS).

Performance of SIS was explored by applying the corresponding weights to the so-called validation set consisting of the 1,048 1:5 case-control sets who did not enter into the training set. To evaluate the clinical utility of SIS for predicting infection, we considered the receiver operating characteristic (ROC) curve analysis and used area under the ROC curve (AUC) as a global summary of the discriminatory capacity of the scores [32].

### Comparing specific and unspecific predictors of SARS-CoV-2 infection

Some unspecific scores surrogating general clinical profile of each case and control included into the study were considered. In particular, the number of drugs with different 3rd level ATC dispensed to, and comorbidities with different ICD-9-CM experienced by each case and control within two-years prior baseline (2018-2019) was recorded. Categorization was made by assigning increasing values of 1, 2, 3 and 4 to 0, 1-4, 5-9 and ≥10 drugs (comedication score) and 1, 2, 3 and 4 to 0, 1-2 and ≥3 comorbidities (comorbidity score). In addition, cases and controls were categorized according to the Multisource Comorbidity Score (MCS), a new index of patients’ clinical status derived from inpatients diagnostic information and outpatient drug prescriptions provided by the regional Italian data and validated for outcome prediction [22,33]. To simplify comparisons, the original five categories of worsening clinical profile (0, 1, 2, 3 and 4) as defined by MCS, were reduced to milder (MCS=0), middle (1≤MCS≤3) and severe (MCS≥4) categories. With the aim of comparing discriminatory ability of specific (SIS) and unspecific (comedications, comorbidities and MCS) predictors of SARS-CoV-2 infection, ROC curves and corresponding AUCs were again used.

All analyses were performed using SAS 9.4 (Cary, NC). A 2-sided p-value of 0.10 or less was considered significant.

## Results

### Clinical predictors of SARS-CoV-2 infection

Table 1 reports univariate and multivariate association between the considered diseases/conditions and the risk of Covid-19 infection. Owing to their low prevalence, fourteen conditions were excluded from this analysis (tuberculosis, weight loss, disorders involving the immune mechanisms, disorders of fluid, electrolyte and acid-base balance, coagulation defects, bipolar disorders, alcohol abuse, drug addiction, multiple sclerosis, cystic fibrosis, chronic and acute pancreatitis, anchylosing spondylitis, systemic sclerosis, systemic sclerosis). Among the 33 remaining conditions, two were grouped, i.e., chronic pulmonary obstructive disease with asthma (chronic respiratory disease), and chronic renal disease with or without dialysis. Among the 31 remaining conditions, 23 (74%) and 12 (39%) showed significant association with the risk of SARS-CoV-2 infection from univariate and multivariable regression respectively (Table 1).

**Table 1.**
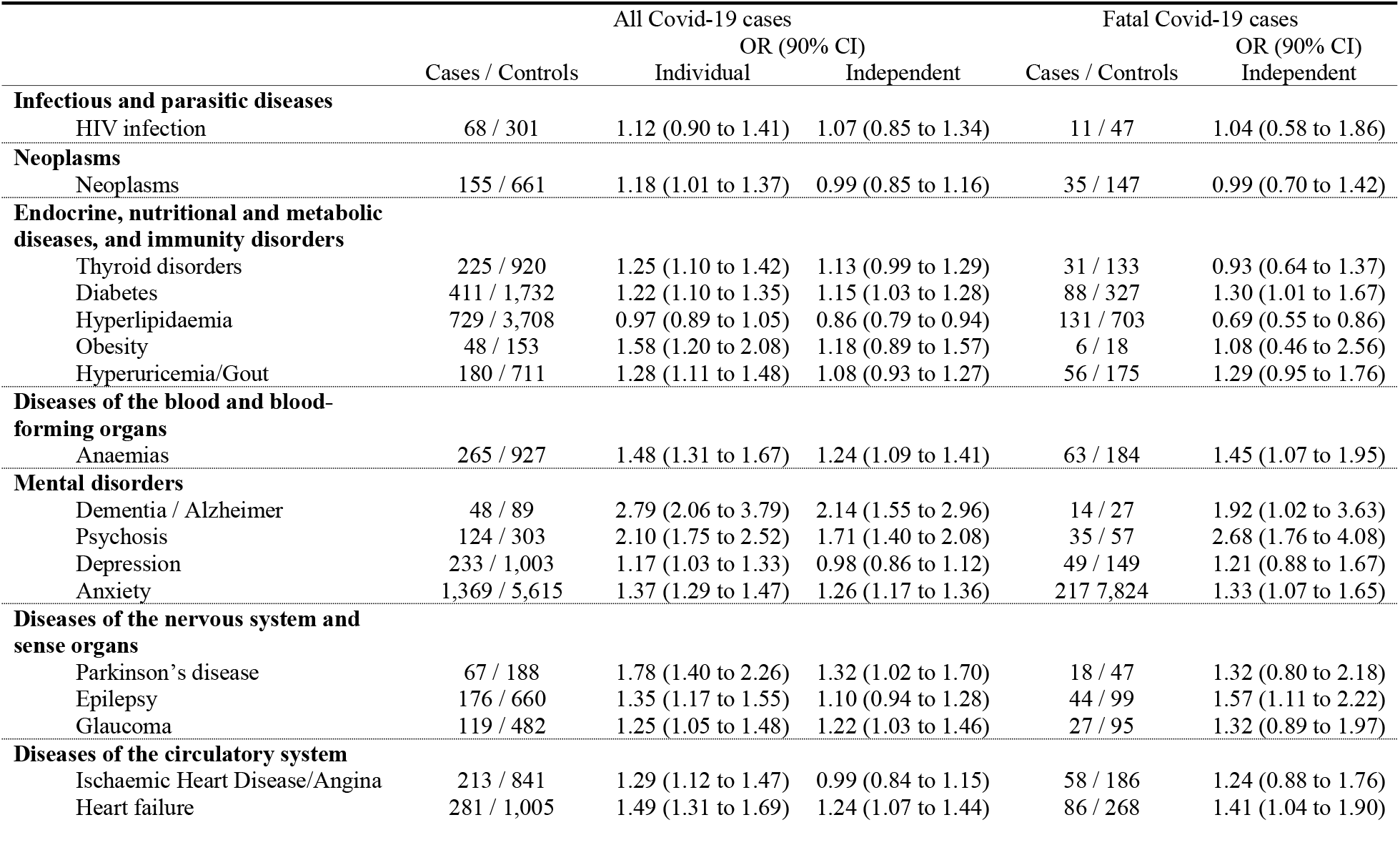
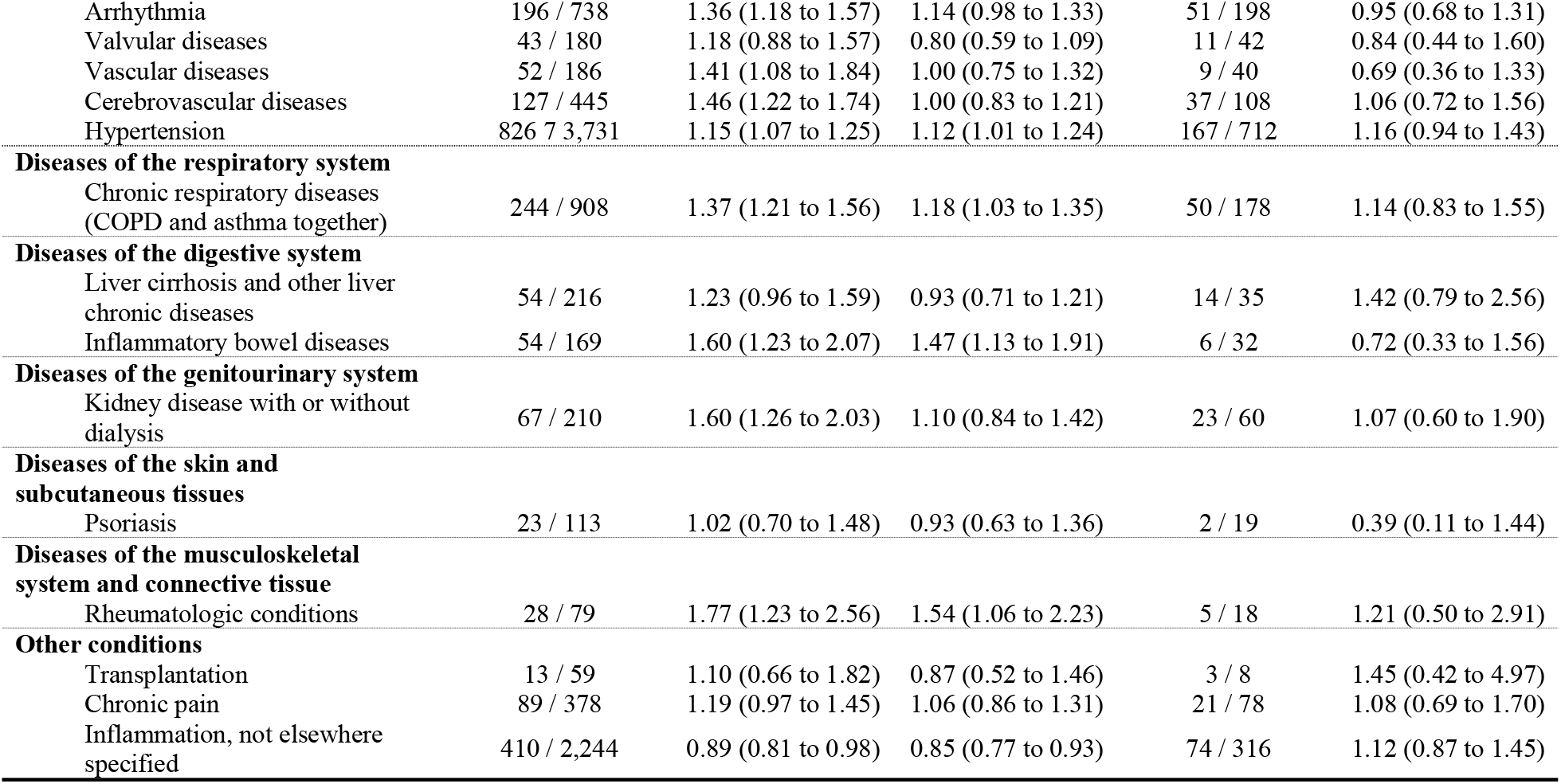
Individual (one by one, univariate) and independent (all together, multivariate) odds ratio (OR), and 90% confidence intervals (CI), for the relationship between selected diseases/conditions and the risk of Covid infection as a whole (3,497 cases and corresponding 17,358 controls), as well as the risk of fatal Covid infection (435 cases and corresponding 2,154 controls).

In particular, patients suffering from diabetes, anaemias, mental disorders (dementia / Alzheimer’s disease, psychosis and anxiety), Parkinson’s disease, glaucoma, diseases of the circulatory system (heart failure and hypertension), chronic respiratory, inflammatory bowel, and rheumatologic conditions showed statistical evidence of increased risk of infection with respect to patients who did not suffer from them. Likely because of low power, only 7 conditions resulted significantly associated with the risk of fatal Covid-19 disease, but there was no relevant difference in the estimates with respect to the risk of SARS-CoV-2 infection as a whole (Table 1).

Same separate analysis was conducted for women and men positive to Covid-19, showing statistical evidence of increased risk infection for women suffering from anaemias, dementia/Alzheimer, psychosis, anxiety, epilepsy, hearth failure, kidney diseases and particularly cystic fibrosis (S3 Table). Otherwise, higher risk of infection was observed among men suffering from diabetes, psychosis, anxiety, Parkinson, arrhythmia, chronic pulmonary disease, inflammatory bowel diseases and particularly dementia/Alzheimer and rheumatologic conditions. Estimates were similar for Covid-19 patients younger and older than 65 years, showing, among the first, significant higher risk of infection for diabetes, anxiety, Parkinson’s disease, arrhythmia, inflammatory bowel and chronic pulmonary diseases, particularly dementia/Alzheimer (S4 Table). While, patients older than 65 years suffering from thyroid disorders, anaemias, dementia/Alzheimer, psychosis, anxiety, epilepsy and hearth failure showed significant higher risk infection.

### SARS-CoV-2 Infection Score (SIS)

Fifteen conditions significantly contributed to the SIS, the corresponding weights being reported in Table 2. Factors which most contributed to the total aggregate score were dementia / Alzheimer’s disease, kidney disease, psychosis, inflammatory bowel disease and rheumatologic conditions, while diabetes, anaemias, anxiety, Parkinson’s disease, glaucoma, heart failure, hypertension, arrhythmia, thyroid disorders and chronic respiratory disease provided small, although significant, contributions. **Fig 1** shows that, as the SIS value increases, the OR progressively increases, being the odds of SARS-CoV-2 infection among people with the highest SIS value (SIS = IV), 1.74 times higher than those unaffected by any SIS contributing conditions (SIS = I). The prevalence of controls stratified according to the SIS score gradually decreases from 50% (SIS = I) to 12% (SIS = IV).

**Table 2.**
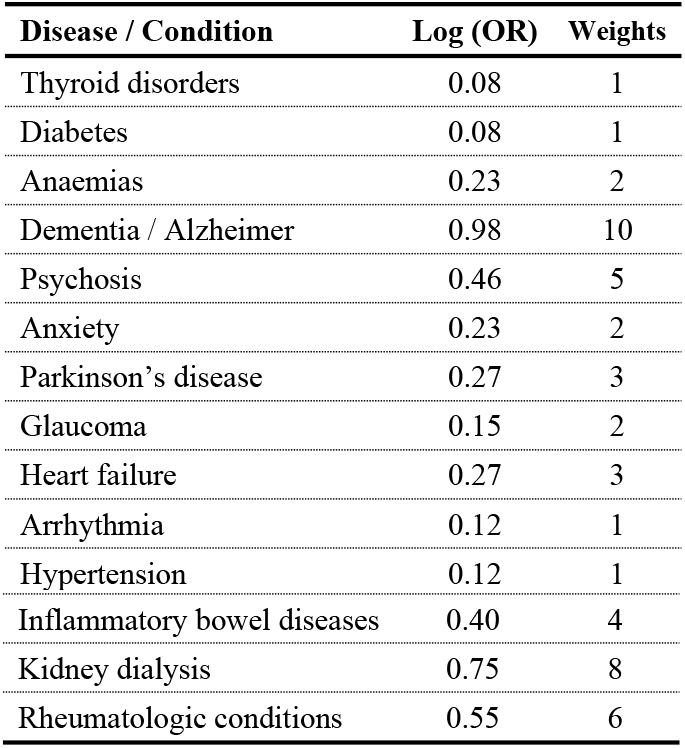
Weights, assigned to diseases that were significantly associated with the risk of Covid-19 disease, used to construct the SARS-CoV-2 Infection Score (SIS).

**Fig 1.**
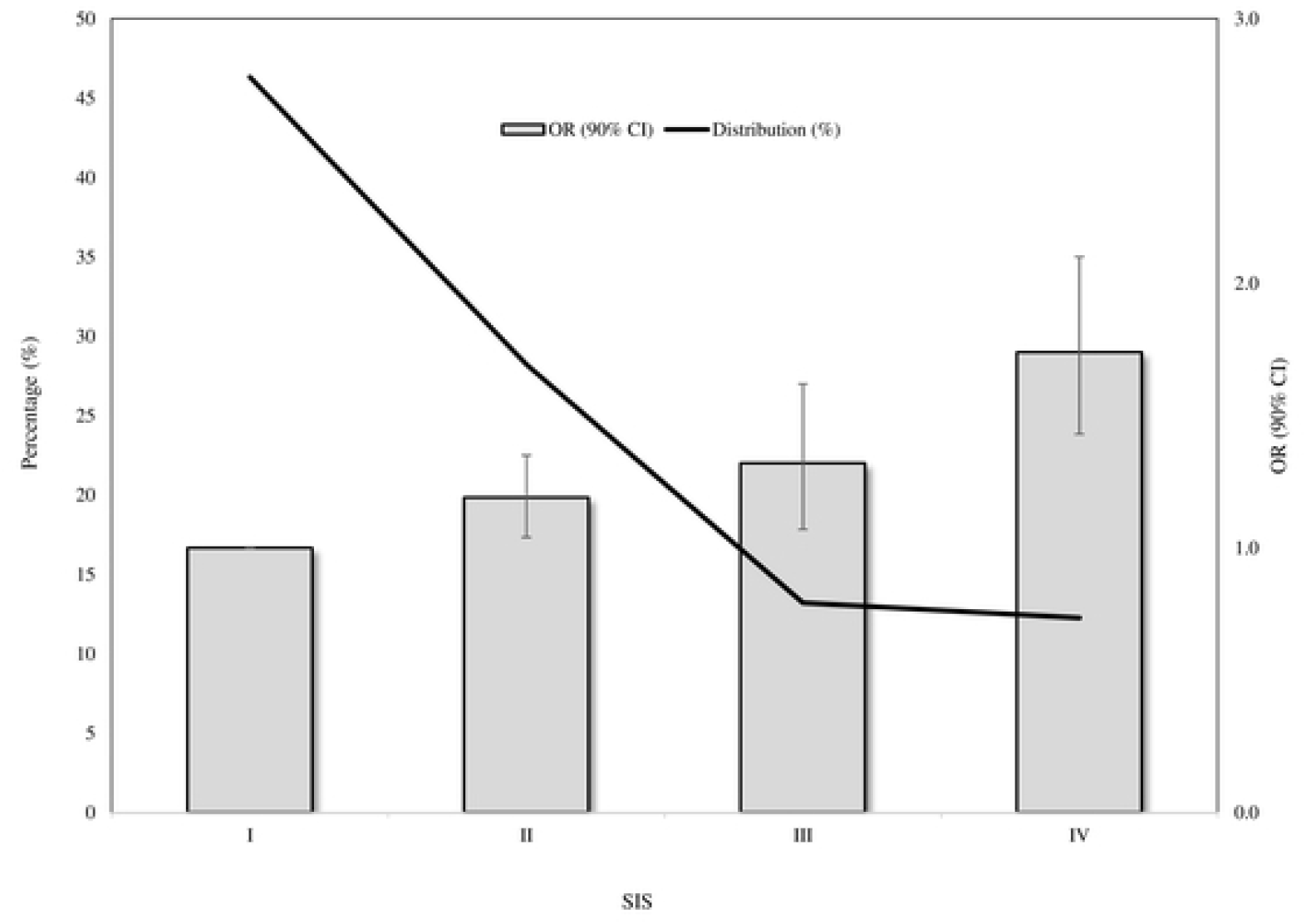
SARS-CoV-2 Infection Score (SIS) distribution among controls, and corresponding trend in odds ratios (and 90% confidence intervals) along categories of SIS. SARS-CoV-2 Infection Score: I, II, III and IV to 0, 1-2, 3-4 and ≥5.

### Comparing with unspecific predictors of SARS-CoV-2 infection

Generic/unspecific scores surrogating clinical profile showed to be associated with the risk of SARS-CoV-2 infection, showing patients with ≥ 10 drug treatments, those with ≥ 3 comorbidities, and those with MCS value ≥ 4, increased risk of 65%, 36% and 45% with respect to patients cotreatments, comorbidities and MCS value = I, respectively (Table 3).

**Table 3.** Relationship between selected score and the risk of SARS-CoV-2 infection.

| Scores | OR (90% CI) |
| --- | --- |
| **SARS-CoV-2 Infection Score (SIS)** |  |
| I (0) | 1.00 (Ref.) |
| II (1-2) | 1.19 (1.03 to 1.36) |
| III (3-4) | 1.32 (1.10 to 1.58) |
| IV ($\geq 5$) | 1.74 (1.44 to 2.10) |
| **Number of comedications** |  |
| I (0) | 1.00 (Ref.) |
| II (1-4) | 1.05 (0.91 to 1.21) |
| III (5-9) | 1.17 (0.97 to 1.41) |
| IV ($\geq 10$) | 1.65 (1.25 to 2.19) |
| **Number of comorbidities** |  |
| I (0) | 1.00 (Ref.) |
| II (1-2) | 1.21 (1.05 to 1.38) |
| III ($\geq 3$) | 1.36 (1.15 to 1.60) |
| **Multisource Comorbidity Score (MCS)** |  |
| I (0) | 1.00 (Ref.) |
| II (1-3) | 1.21 (1.03 to 1.41) |
| III ($\geq 4$) | 1.45 (1.23 to 1.70) |

AUC (90% CI) of SIS, cotreatment and comorbidity scores and MCS respectively had values of 0.54 (0.52 to 0.56), 0.52 (0.50 to 0.54), 0.53 (0.51 to 0.55), and 0.53 (0.51 to 0.55) (Fig 2). There was no evidence that specific and unspecific scores had different discriminatory ability.

**Fig 2.**
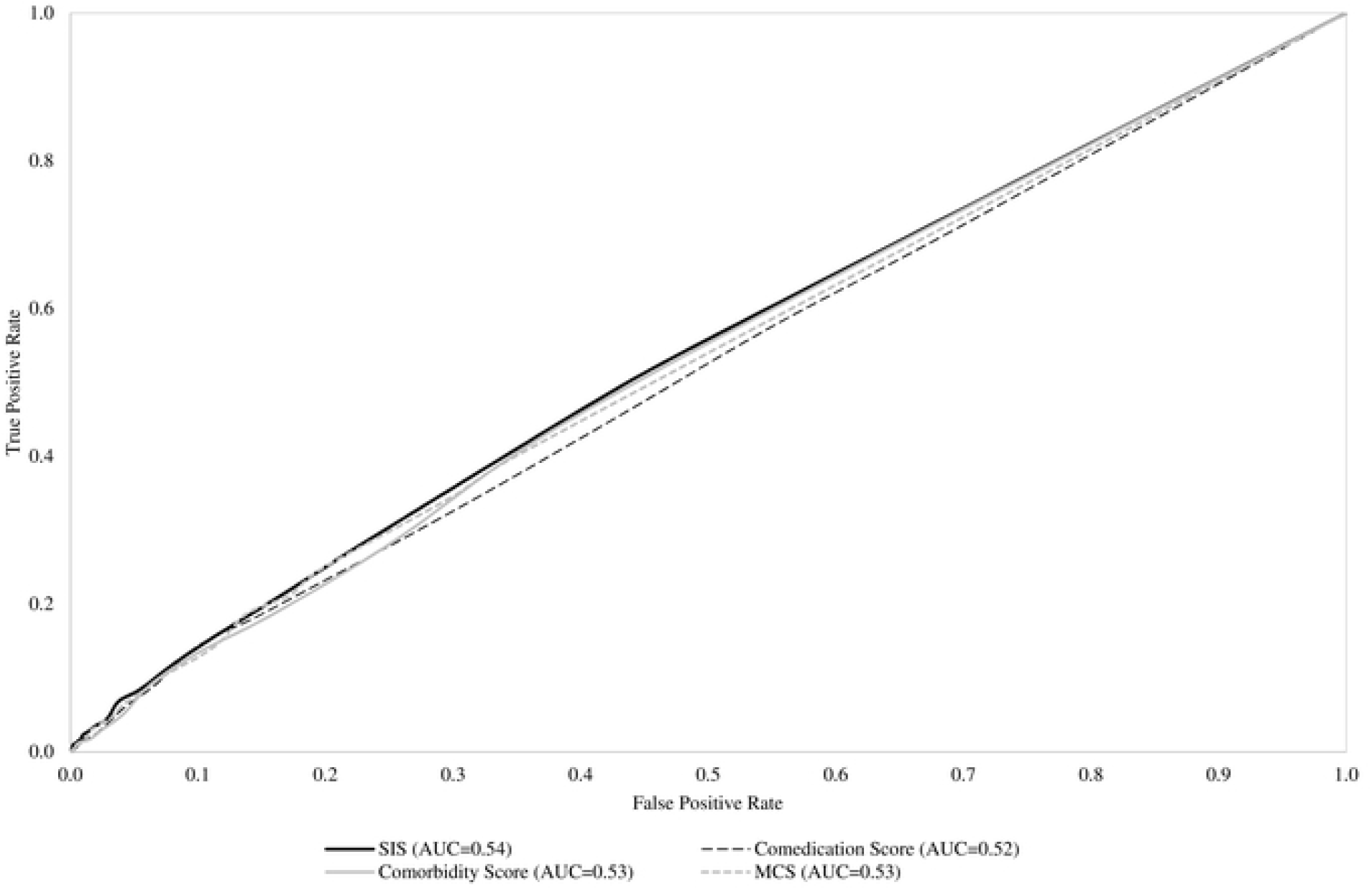
Receiver Operating Characteristics (ROC) curves comparing discriminant power of SARS-CoV-2 Infection Score (SIS), and selected unspecific score surrogating clinical profile (cotreatments, comorbidities and Multisource Comorbidity Score).

## Discussion

Our study shows that several diseases and conditions are significantly and independently associated with the risk of SARS-CoV-2 infection. Beyond conditions making particularly vulnerable the respiratory system (e.g., chronic obstructive pulmonary disease and asthma), comorbidities including practically all diagnostic categories are involved. Predictors belonging to nutritional and metabolic (diabetes), cardiovascular (heart failure and hypertension) and renal diseases were widely expected, since it has accepted that SARS-CoV-2 has major implications for the cardiovascular system. Indeed, patients with heart failure [34], diabetes [35–37], hypertension [38] and kidney disease [39–41] have been consistently identified as particularly vulnerable populations, and these findings were consistently found in our study. In addition, we confirmed that people with weakened immune systems from a medical condition or treatment are at a higher risk. Among these, those living with haemoglobin disorders [42], inflammatory bowel disease [43] and immune-rheumatological diseases [44] must be considered vulnerable groups for Covid-19 infection. Mental health and cognitive function might have independent utility in understanding the burden of respiratory disease, since they may influence the risk of contracting the infection, at least in part by impairing innate or adaptive immunity [45] and diminishing the precautions taken to minimize risk. Another explanation of our findings is that people with history of depression [46], psychosis [47] and stress disorders [48] could experience elevated rates of an array of respiratory infections because these conditions often require treatment in a psychiatric care facility, and the risk of infection can be particularly high in these structures. Finally, our study adds evidence regarding the impact of diseases and conditions on the risk of SARS-CoV-2 infection between men and women. As pointed out by a recent study [49], sex and age disaggregated data are essential for understanding the distributions of risk infection in the population and the extent to which they affect clinical outcomes.

Despite our results confirm that a wide range of diseases and conditions likely increase vulnerability to SARS-CoV-2 infection, and probably its more severe clinical manifestations, we have not been able to develop a score that accurately may predict the risk of infection. In addition, we found that predictive ability of the score obtained by weighting risk factors of SARS-CoV-2 infection, was not better than generic scores of comorbidity and comedication. This expands upon previous findings of individual comorbidities as independent risk factors for SARS-CoV-2 infection [50,51], and confirms our substantial inability to predict the risk of SARS-CoV-2 infection. The reasons are likely linked with the several limitations of our approach that, in general, generates estimates biased towards the null. First, exposure misclassification regards our inability to careful capturing conditions and diseases through algorithms based on healthcare utilization databases [52]. Second, it is well known that outcome misclassification can bias epidemiologic results. For Covid-19, suboptimal test sensitivity, despite excellent specificity, results in an overestimation of cases in the early stages of an outbreak, and substantial underestimation of cases as prevalence increases [53]. It should be noticed, however, that both, exposure and outcome misclassification likely drew estimates towards the null (i.e., underestimate the strength of the association between their presence and the outcome risk) so generating uncertainty for the weighting approach of score developing. Third, the lack of specific data regarding the clinical outcome for the stratification of Covid-19 positive patients in terms of home isolation, hospitalization and admission in intensive care. Fourth, the lack of information on biologic markers potentially able to predict infection, and severity of its clinical manifestations, is another limitation of our study, as for example, according to the current literature, some laboratory hallmarks have been shown to predict infection, particularly in more severe cases [54]. Finally, our choice of accepting a 0.10 first type error, and of consequently reporting 90% confidence intervals, is justified by the exploratory nature of our study, but at the same time likely generate false positive signals, so limiting discriminant power of the score.

In conclusion, taking the limitations we discussed into account, we identified conditions and diseases that make people more vulnerable to SARS-CoV-2 infection. These findings contribute to inform public health, and clinical decisions regarding risk stratifying. However, further research is need for developing a score reliably predicting the risk, possibly by integrating healthcare utilization with clinical and biological data.

Our results can be an important tool supporting all clinical and political stakeholders allowing the identification of the population most at risk of contracting Covid-19 and facilitating the provision of appropriate preventive/therapeutic measures, especially with the hypothetic prediction of a new autumn outbreak. Adopting preventive measures can help to minimize the damage generated by a potential new relapse that the health systems will face.

## Supporting Information

**S1 Table. Campania Region Database (CaReDB) characteristics.** ATC = Anatomical Therapeutic Chemical; ICD-9-CM = *International Classification of Diseases*, 9th Revision, Clinical Modification. ^a^Time span covered was 2009–2018 for hospital-discharge records and 2014–2019 for outpatient pharmacy records.

**S2 Table. List of diseases and conditions candidate for predicting SARS-CoV-2 infection, and corresponding ICD-CM and ATC codes used for detecting they.**

**S3 Table. Odds ratio (OR), and 90% confidence intervals (CI), for the relationship between selected diseases/conditions and the risk of SARS-CoV-2 infection, stratified according to gender.**

**S4 Table. Odds ratio (OR), and 90% confidence intervals (CI), for the relationship between selected diseases/conditions and the risk of SARS-CoV-2 infection, stratified according to age categories (i.e., younger and older 65 years)**

## References

1. Dong Ensheng, Du Hongru, Gardner Lauren. An interactive web-based dashboard to track COVID-19 in real time. The Lancet Infectious Diseases 2020;20:533–4.

2. Onder G, Rezza G, Brusaferro S. Case-Fatality Rate and Characteristics of Patients Dying in Relation to COVID-19 in Italy [published online ahead of print, 2020 Mar 23]. JAMA. 2020;10.1001/jama.2020.4683. doi:10.1001/jama.2020.4683.

3. Liang W, Liang H, Ou L, Chen B, Chen A, Li C, et al. Development and Validation of a Clinical Risk Score to Predict the Occurrence of Critical Illness in Hospitalized Patients With COVID-19. JAMA Intern Med. 2020;e202033. doi:10.1001/jamainternmed.2020.2033.

4. Xie J, Hungerford D, Hui Chen H, Abrams ST, Li S, Wang G, et al. Development and external validation of a prognostic multivariable model on admission for hospitalized patients with COVID-19. medRxiv preprint doi:10.1101/2020.03.28.20045997.

5. Li Q, Zhang J, Ling Y, Li W, Zhang X, Luet H, al. A simple algorithm helps early identification of SARS-CoV-2 infection patients with severe progression. Infection. 2020; 1–8. doi:10.1007/s15010-020-01446-z.

6. Sun Y, Koh V, Marimuthu K, Tek Ng O, Young B, Vasoo S, et al. Epidemiological and Clinical Predictors of COVID-19. Clin Infect Dis. 2020;ciaa322. doi:10.1093/cid/ciaa322.

7. Haimovich A, Ravindra NG, Stoytchev S, Young HP, Wilson FP, van Dijk D, et al. Development and validation of the COVID-19 severity index (CSI): a prognostic tool for early respiratory decompensation. medRxiv preprint. doi:10.1101/2020.05.07.20094573.

8. Zheng Z, Peng F, Xu B, Zhao J, Liu H, Peng J, et al. Risk factors of critical & mortal COVID-19 cases: A systematic literature review and meta-analysis. J Infect. 2020; S0163-4453(20) 30234–6. doi:10.1016/j.jinf.2020.04.021.

9. Moreno Juste A, Menditto E, Orlando V, Monetti VM, Gimeno Miguel A, González Rubio F, et al. Treatment Patterns of Diabetes in Italy: A Population-Based Study. Front Pharmacol. 2019;10:870. Published 2019 Aug 6. doi:10.3389/fphar.2019.00870.

10. Guerriero F, Orlando V, Monetti VM, Russo V, Menditto E. Biological therapy utilization, switching, and cost among patients with psoriasis: retrospective analysis of administrative databases in Southern Italy. Clinicoecon Outcomes Res. 2017;9:741–748. Published 2017 Dec 1. doi:10.2147/CEOR.S147558.

11. Russo V, Monetti VM, Guerriero F, Trama U, Guida A, Menditto E, et al. Prevalence of antibiotic prescription in southern Italian outpatients: real-world data analysis of socioeconomic and sociodemographic variables at a municipality level. Clinicoecon Outcomes Res. 2018;10:251–258. Published 2018 May 3. doi:10.2147/CEOR.S161299.

12. Iolascon G, Gimigliano F, Orlando V, Capaldo A, Di Somma C, Menditto E. Osteoporosis drugs in real-world clinical practice: an analysis of persistence. Aging Clin Exp Res. 2013;25 Suppl 1:S137–S141. doi:10.1007/s40520-013-0127-5.

13. Orlando V, Guerriero F, Putignano D, Monetti VM, Tari DU, Farina G, et al. Prescription Patterns of Antidiabetic Treatment in the Elderly. Results from Southern Italy. Curr Diabetes Rev. 2015;12(2):100–106. doi:10.2174/1573399811666150701120408.

14. Menditto E, Cahir C, Aza-Pascual-Salcedo M, Bruzzese D, Poblador-Plou B, Malo S, et al. Adherence to chronic medication in older populations: application of a common protocol among three European cohorts. Patient Prefer Adherence. 2018; 12:1975–1987. Published 2018 Oct 5. doi:10.2147/PPA.S164819.

15. Casula M, Catapano AL, Piccinelli R, Menditto E, Manzoli L, De Fendi L, et al. Assessment and potential determinants of compliance and persistence to antiosteoporosis therapy in Italy. Am J Manag Care. 2014;20(5):e138–e145.

16. Orlando V, Monetti VM, Moreno Juste A, Russo V, Mucherino S, Trama U, et al. Drug Utilization Pattern of Antibiotics: The Role of Age, Sex and Municipalities in Determining Variation. Risk Manag Healthc Policy. 2020; 13:63–71. Published 2020 Jan 29. doi:10.2147/RMHP.S223042.

17. Orlando V, Coscioni E, Guarino I, Mucherino S, Perrella A, Trama U, et al. Drug-utilisation Profiles and COVID-19: Retrospective Cohort Study in Italy, 29 May 2020, PREPRINT (Version 1) available at Research Square. doi: 10.21203/rs.3.rs-31829/v.

18. Corman VM, Landt O, Kaiser M, Molenkamp R, Meijer A, Chu DKW, et al. Detection of 2019 novel coronavirus (2019-nCoV) by 9 real-time RT-PCR. Euro Surveill. 2020 Jan;25(3):2000045. doi: 10.2807/1560-107917.ES.2020.25.3.2000045.

19. Italian Data Protection Authority. General authorisation to process personal data for scientific research purposes – 1 March 2012 [1884019]. 10.1094/PDIS-11-11-0999-PDN.

20. O’Shea M, Teeling M, Bennett K. The prevalence and ingredient cost of chronic comorbidity in the Irish elderly population with medication treated type 2 diabetes: A retrospective crosssectional study using a national pharmacy claims database. BMC Health Serv Res 2013;13(1):1. doi: 10.1186/1472-6963-13-23.

21. Pratt NL, Kerr M, Barratt JD, Kemp-Casey A, Kalisch Ellett LM, Ramsay E, et al. The validity of the Rx-Risk Comorbidity Index using medicines mapped to the Anatomical Therapeutic Chemical (ATC) Classification System. BMJ Open 2018;8(4):1–8. doi: 10.1136/bmjopen-2017-021122.

22. Corrao G, Rea F, Di Martino M, De Palma R, Scondotto S, Fusco D, et al. Developing and validating a novel multisource comorbidity score from administrative data: a large populationbased cohort study from Italy. BMJ Open. 2017;7(12):e019503. Published 2017 Dec 26. doi:10.1136/bmjopen-2017-019503.

23. Leebeek FW, Eikenboom JC. Von Willebrand’s Disease. N Engl J Med. 2016;375(21):2067–2080. doi:10.1056/NEJMra1601561.

24. Maio V, Yuen E, Rabinowitz C, Louis D, Jimbo M, Donatini A, et al. Using pharmacy data to identify those with chronic conditions in Emilia Romagna, Italy. Journal of health services research & policy, 2005, 10.4: 232–238. doi:10.1258/135581905774414259.

25. Aboujaoude E, Salame WO. Naltrexone: A Pan-Addiction Treatment? CNS Drugs. 2016;30(8):719–733. doi:10.1007/s40263-016-0373-0.

26. Canova C, Danieli S, Barbiellini CA, Simonato L, Di RD, Cappai G, et al. A Systematic Review of Case-Identification Algorithms Based on Italian Healthcare Administrative Databases for Three Relevant Diseases of the Nervous System: Parkinson’s Disease, Multiple Sclerosis, and Epilepsy. Epidemiol Prev. 2019;43(4 Suppl 2):62–74. doi:10.19191/EP19.4.S2.P062.093.

27. Bezzini D, Policardo L, Meucci G, Ulivelli M, Bartalini S, Profili F, et al. Prevalence of Multiple Sclerosis in Tuscany (Central Italy): A Study Based on Validated Administrative Data. Neuroepidemiology. 2016;46(1):37–42. doi:10.1159/000441567.

28. Bargagli AM, Colais P, Agabiti N, Mayer F, Buttari F, Centonze D, et al. Prevalence of multiple sclerosis in the Lazio region, Italy: use of an algorithm based on health information systems. J Neurol. 2016;263(4):751–759. doi:10.1007/s00415-016-8049-8.

29. Chini F, Pezzotti P, Orzella L, Borgia P, Guasticchi G. Can we use the pharmacy data to estimate the prevalence of chronic conditions? a comparison of multiple data sources. BMC Public Health. 2011;11:688. Published 2011 Sep 5. doi:10.1186/1471-2458-11-688.

30. Tibshirani R. The lasso method for variable selection in the Cox model. Stat Med. 1997;16(4):385–395. doi:10.1002/(sici)1097-0258(19970228)16:4<385::aid-sim380>3.0.co;2-3.

31. Gagne JJ, Glynn RJ, Avorn J, Levin R, Schneeweiss S. A combined comorbidity score predicted mortality in elderly patients better than existing scores. J Clin Epidemiol. 2011;64(7):749–759. doi:10.1016/j.jclinepi.2010.10.004.

32. Xu H, Qian J, Paynter NP, Zhang X, Whitcomb BW, Tworoger SS, et al. Estimating the receiver operating characteristic curve in matched case control studies. Stat Med. 2019;38(3):437–451. doi:10.1002/sim.7986.

33. Corrao G, Rea F, Carle F, Di Martino M, De Palma R, Francesconi P, et al. Measuring multimorbidity inequality across Italy through the multisource comorbidity score: a nationwide study [published online ahead of print, 2020 May 20]. Eur J Public Health. 2020;ckaa063. doi:10.1093/eurpub/ckaa063.

34. Wang BX. Susceptibility and prognosis of COVID-19 patients with cardiovascular disease. Open Heart. 2020;7(1):e001310. doi:10.1136/openhrt-2020-001310.

35. Carey IM, Critchley JA, DeWilde S, Harris T, Hosking FJ, Cook DG. Risk of Infection in Type 1 and Type 2 Diabetes Compared With the General Population: A Matched Cohort Study. Diabetes Care. 2018;41(3):513–521. doi:10.2337/dc17-2131.

36. Geerlings SE, Hoepelman AI. Immune dysfunction in patients with diabetes mellitus (DM). FEMS Immunol Med Microbiol. 1999;26(3-4):259–265. doi:10.1111/j.1574-695X.1999.tb01397.x.

37. Peleg AY, Weerarathna T, McCarthy JS, Davis TM. Common infections in diabetes: pathogenesis, management and relationship to glycaemic control. Diabetes Metab Res Rev. 2007;23(1):3–13. doi:10.1002/dmrr.682.

38. Mancia G, Rea F, Ludergnani M, Apolone G, Corrao G. Renin-Angiotensin-Aldosterone System Blockers and the Risk of Covid-19. N Engl J Med. 2020;382(25):2431–2440. doi:10.1056/NEJMoa2006923.

39. Perico L, Benigni A, Remuzzi G. Should COVID-19 Concern Nephrologists? Why and to What Extent? The Emerging Impasse of Angiotensin Blockade. Nephron. 2020;144(5):213–221. doi:10.1159/000507305.

40. Cheng Y, Luo R, Wang K, Zhang M, Wang Z, Dong L et al. Kidney disease is associated with in-hospital death of patients with COVID-19. Kidney Int. 2020;97(5):829–838. doi:10.1016/j.kint.2020.03.005

41. Diao B, Feng Z, Wang C, Feng Z, Tan Y, Wang H, et al. Human kidney is a target for novel severe acute respiratory syndrome coronavirus 2 (SARS-CoV-2) infection. medRxiv. In press. doi: 10.1101/2020.03.04.20031120.

42. Chowdhury SF, Anwar S. Management of Hemoglobin Disorders During the COVID-19 Pandemic. Front Med (Lausanne). 2020;7:306. Published 2020 Jun 9. doi:10.3389/fmed.2020.00306.

43. de León-Rendón JL, Hurtado-Salazar C, Yamamoto-Furusho JK. Aspects of inflammatory bowel disease during the COVID-19 pandemic and general considerations. Aspectos y consideraciones generales en la enfermedad inflamatoria intestinal durante la pandemia por COVID-19. Rev Gastroenterol Mex. 2020;S0375-0906(20)30054-9. doi:10.1016/j.rgmx.2020.05.001.

44. Favalli EG, Ingegnoli F, De Lucia O, Cincinelli G, Cimaz R, Caporali R. COVID-19 infection and rheumatoid arthritis: Faraway, so close!. Autoimmun Rev. 2020;19(5):102523. doi:10.1016/j.autrev.2020.102523.

45. Cohen S, Tyrrell DA, Smith AP. Psychological stress and susceptibility to the common cold. N Engl J Med. 1991;325(9):606–612. doi:10.1056/NEJM199108293250903.

46. Andersson NW, Goodwin RD, Okkels N, Gustafsson LN, Taha F, Cole SW et al. Depression and the risk of severe infections: prospective analyses on a nationwide representative sample. Int J Epidemiol. 2016;45(1):131–139. doi:10.1093/ije/dyv333.

47. Seminog OO, Goldacre MJ. Risk of pneumonia and pneumococcal disease in people with severe mental illness: English record linkage studies. Thorax. 2013;68(2):171–176. doi:10.1136/thoraxjnl-2012-202480.

48. Jiang T, Farkas DK, Ahern TP, Lash TL, Sørensen HT, Gradus JL. Posttraumatic Stress Disorder and Incident Infections: A Nationwide Cohort Study. Epidemiology. 2019;30(6):911–917. doi:10.1097/EDE.0000000000001071.

49. Sharma G, Volgman AS, Michos ED. Sex Differences in Mortality from COVID-19 Pandemic: Are Men Vulnerable and Women Protected? JACC Case Rep. 2020;10.1016/j.jaccas.2020.04.027. doi:10.1016/j.jaccas.2020.04.027.

50. Guan WJ, Liang WH, Zhao Y, Liang H, Chen Z, Li Y et al. Comorbidity and its impact on 1590 patients with COVID-19 in China: a nationwide analysis. Eur Respir J. 2020;55(5):2000547. Published 2020 May 14. doi:10.1183/13993003.00547-2020.

51. Christensen DM, Strange JE, Gislason G, Torp-Pedersen C, Gerds T, Fosbøl E et al. Charlson Comorbidity Index Score and Risk of Severe Outcome and Death in Danish COVID-19 Patients. J Gen Intern Med. 2020;1–3. doi:10.1007/s11606-020-05991-z.

52. Burstyn I, Yang Y, Schnatter AR. Effects of non-differential exposure misclassification on false conclusions in hypothesis-generating studies. Int J Environ Res Public Health. 2014;11(10):10951–10966. Published 2014 Oct 21. doi:10.3390/ijerph111010951.

53. Burstyn I, Goldstein ND, Gustafson P. Towards reduction in bias in epidemic curves due to outcome misclassification through Bayesian analysis of time-series of laboratory test results: Case study of COVID-19 in Alberta, Canada and Philadelphia, USA. Preprint. medRxiv. 2020;2020.04.08.20057661. Published 2020 Apr 11. doi:10.1101/2020.04.08.20057661.

54. Vultaggio A, Vivarelli E, Virgili G, Lucenteforte E, Bartoloni A, Nozzoli C et al. Prompt predicting of early clinical deterioration of moderate-to-severe COVID-19 patients: usefulness of a combined score using IL-6 in a preliminary study. J Allergy Clin Immunol Pract 2020:S2213-2198(20)30611-5.

